# Nanoscale regulation of ROS signaling at the plasma membrane tunes the plant response to osmotic stress

**DOI:** 10.64898/2026.01.19.698185

**Authors:** Arthur Poitout, José M. Ugalde, Lucille Gorgues, Armelle Dongois, Patricia Scholz, Philippe Nacry, Carine Alcon, Jean-Bernard Fiche, Xavier Dumont, Marcelo Nollman, Komal Jhala, Anton Schäffner, Yvon Jaillais, Lionel Verdoucq, Andreas J. Meyer, Alexandre Martinière

## Abstract

The spatiotemporal organization of proteins and lipids within membranes is crucial for ensuring proper cellular signaling. While the segregation of proteins and lipids into membrane nanodomains is well established, it remains unclear whether nanodomains can generate gradients of small diffusible molecules. In plants, reactive oxygen species (ROS), especially hydrogen peroxide (H_2_O_2_), act as a key signaling molecules in response to environmental stimuli such as osmotic stress. However, how extracellular H_2_O_2_ affects intracellular signaling has remained unknown. Here, we show that osmotic stimulation induces the formation of localized, H_2_O_2_-rich nano-environments at the cytoplasmic face of the plasma membrane (PM) in *Arabidopsis* root cells. Using a PM-tethered H_2_O_2_ biosensor, we found that these oxidized nanodomains arise from the clustering of RESPIRATORY BURST OXIDASE HOMOLOGs (RBOHs) and RHO OF PLANTS 6 (ROP6), in coordination with aquaporin-mediated H_2_O_2_ transport via the PLASMA MEMBRANE INTRINSIC PROTEIN2;7 (PIP2;7). These local redox hotspots at the PM create a feedforward loop in which H_2_O_2_ enhances ROP6 nanoclustering thereby amplifying ROS signaling. Disruption of H_2_O_2_ production or transport dampens both ROP6 clustering and anisotropic cell expansion, indicating a crucial role for spatially-confined redox signaling in regulating plant growth under osmotic stress. Our findings propose a model in which ROP6/RBOHD-F/PIP2;7 nanodomains function as discrete redox signaling units, redefining ROS signaling at the PM as a structured, signal-specific, and compartmentalized process.

## INTRODUCTION

The plasma membrane (PM) is a dynamic mosaic of lipids and proteins organized into heterogeneous patches of distinct size, composition and function, commonly referred to as nanodomains (1–3). These nanodomains are recognized as platforms modulating protein-protein and protein-lipid interactions, thereby playing central roles in the spatial and temporal regulation of signaling pathways. As such, biological membranes are not passive barriers but active participants in signal integration (1). A major challenge in cell biology is to elucidate how membrane organization and dynamics quantitatively influence core features of signaling such as specificity, processing, and amplification, ultimately shaping cellular responses.

In plants, the production of reactive oxygen species (ROS) triggers a broad set of molecular responses to environmental stimuli as well as key developmental processes. In many cases, ROS generation is initiated in the extracellular matrix (the apoplast) where superoxide (O_2_^-^) is produced by NADPH oxidases, known as RESPIRATORY BURST OXIDASE HOMOLOGs (RBOHs) in plants (4–6). Superoxide is rapidly converted into hydrogen peroxide (H_2_O_2_), which serves as a more stable signaling molecule.

Apoplastic H_2_O_2_ can be directly perceived by PM-localized receptor-like kinases such as HYDROGEN-PEROXIDE-INDUCED Ca²⁺ INCREASES 1 (HPCA1), which triggers cytosolic Ca²⁺ influx and contributes to the propagation of systemic ROS waves for long-distance signaling (7–9). In parallel, a fraction of apoplastic H_2_O_2_ diffuses across the PM and reach the cytosol, where it can oxidize thiol groups to sulfenic acid. These intermediates may further react to form mixed disulfides with glutathione or other proteins, or form intramolecular disulfide bonds (10, 11). Such post-translational modifications (PTMs) often modulate the localization, conformation, or activity of target proteins (12–14).

Despite advances in our understanding of the perception and integration of ROS signals, the spatial regulation of H_2_O_2_-dependent signaling has remained poorly understood. For instance, it remains unclear whether elevated H_2_O_2_ levels are uniform throughout the cell or whether variations in H_2_O_2_ levels exist between subcellular regions during signaling events. Addressing this gap is key to understand the specificity and dynamics of ROS-mediated signaling.

Rho of Plants (ROPs) form a plant-specific clade of small GTPases that play essential roles in regulating polarized cell growth, such as in root hairs and pollen tubes (15–17). Like other Rho-family GTPases, ROPs function as molecular switches that cycle between an inactive GDP-bound form and an active GTP-bound state. In their active conformation, ROPs interact with a range of effector proteins to modulate diverse downstream cellular processes (15). Notably, direct physical interactions between GTP-bound ROPs and RBOHs have been demonstrated through both, biochemical and structural approaches (18–22). These interactions suggest a mechanistic link between ROP activation and ROS production. Genetic evidence further supports this connection given that overexpression of constitutively active (CA) ROPs leads to elevated ROS levels, while ROP loss-of-function mutants exhibit impaired ROS accumulation (23–26). Together, these findings suggest that ROPs are positive regulators of RBOH-mediated ROS production, integrating membrane-localized signaling cues with redox-based cellular responses.

Previous studies have shown that ROP6 bind to the PM in root epidermal cells via prenylation of its C-terminus (25, 27). Using single-molecule localization microscopy, it was shown that within minutes of hyperosmotic stimulation, a subset of mEOS2-ROP6 molecules reorganizes into nanodomains of approximately 50 nm in diameter within the PM (21, 28). These ROP6 nanodomains are critical signaling hubs, required for the ROS increase in response to osmotic stress. Importantly, osmotic stimulation leads to the co-localization of ROP6 with RBOHD and RBOHF within the nanodomains. This suggests that osmotic cues trigger the activation and spatial reorganization of ROP6, promoting its clustering with RBOHD/F to initiate ROS production. However, while these observations strongly support a role for ROP6/RBOH nanodomains in ROS signaling, it remains unresolved if these clusters constitute a functional and spatially regulated signaling unit that integrates ROS production with signal specificity at the PM during osmotic stress.

Using the genetically encoded H_2_O_2_–sensitive biosensor HyPer7 (29, 30), this study reveals that osmotic stimulation induces the formation of H_2_O_2_-enriched nano-environments on the cytoplasmic side of the PM. These oxidized nanodomains arise from the localized activity of RBOHs and a facilitated H_2_O_2_ diffusion, specifically within ROP6-containing nanodomains. Surprisingly, the spatially restricted oxidation at the nanodomains creates a feedforward loop, in which the local increase of H_2_O_2_ levels further promotes ROP6 clustering. This, in turn, amplifies ROS production and contributes to cellular growth under osmotic stress.

## RESULTS

### ROP6-HyPer7 allows detection of osmotically-induced ROS in *Arabidposis* root cells

To monitor osmotically-induced H_2_O_2_ accumulation near its production site, we generated a fusion construct between HyPer7, a highly sensitive hydrogen peroxide (H_2_O_2_) ratiometric sensor (29, 30) and ROP6 that interacts with the NADPH oxidase RBOHD/F after osmotic stimulation (18–22) (Figure 1A). The HyPer7 protein was fused to the N-terminus of ROP6 to avoid interfering with its function (21, 27, 31). As expected, the HyPer7-ROP6 signal in Arabidopsis root tip cells localized predominantly at the PM, with only minimal cytosolic labeling, compared to the diffuse distribution observed with soluble HyPer7 (Figure 1B). To assess the fluorescent properties of HyPer7-ROP6 *in planta*, we treated seedlings with increasing concentrations of H_2_O_2_ and compared their responses to plants expressing soluble HyPer7. The response curves of both lines were similar, indicating that the ROP6 fusion does not compromise the dynamic range of the HyPer7 sensor (Fig. S1 A). Furthermore, plants expressing either HyPer7 or HyPer7-ROP6 displayed no significant growth defects compared to the Col-0 wild type (Fig. S1 B).

**Figure 1:**
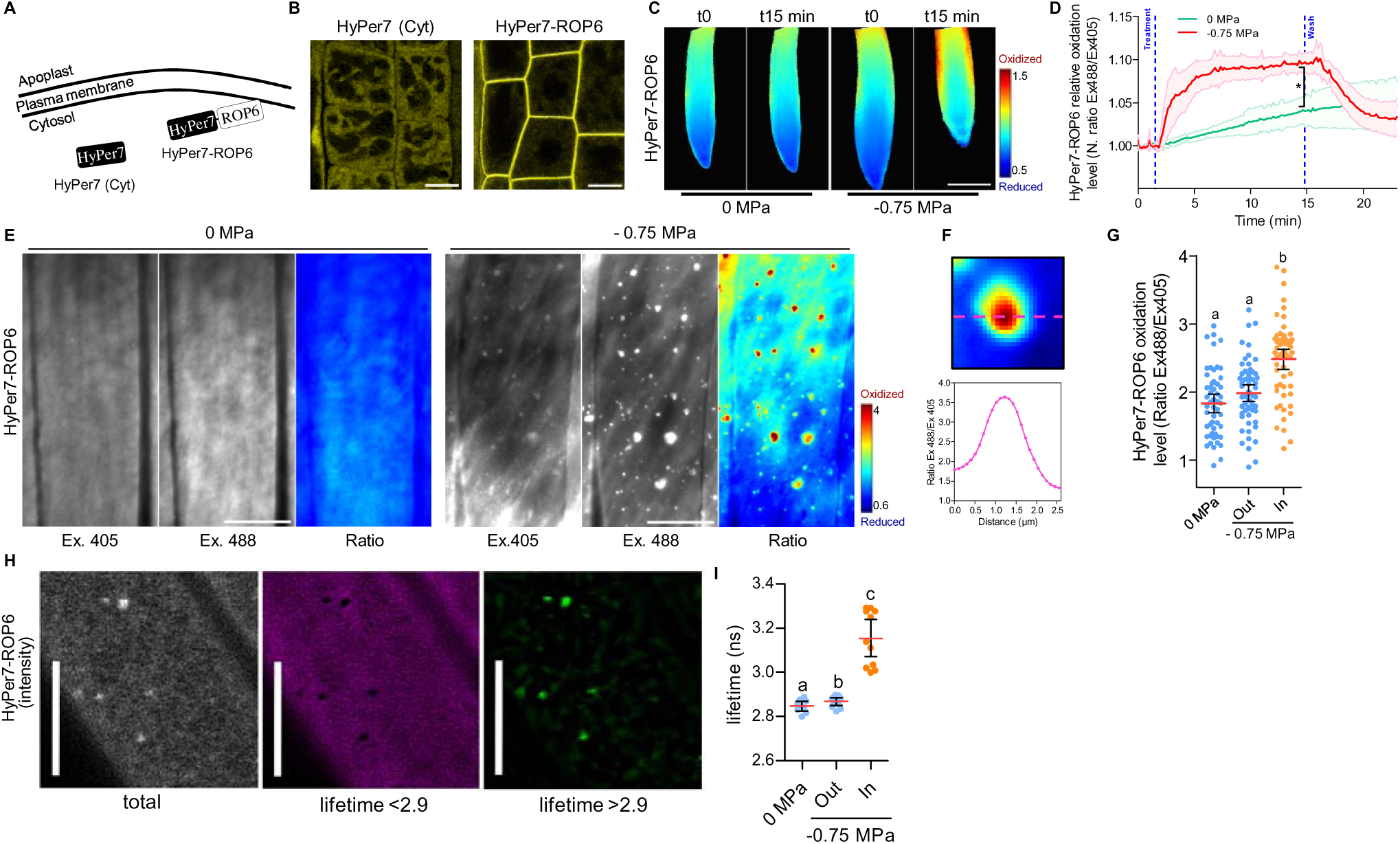
HyPer7-ROP6 reveals formation of ROS nanoenvironments at the PM in response to osmotic shock A, Scheme of the cellular localization of HyPer7 sensor in its free form or fused to ROP6. B, Confocal images of epidermal root tip cells expressing HyPer7 or HyPer7-ROP6. C, Ratiometric images of 5 days old Arabidopsis plants expressing HyPer7-ROP6 in control condition (0 MPa) or upon osmotic stimulation (-0.75 MPa). D, Typical time course showing HyPer7-ROP6 oxidation level (N. Ratio Ex488/Ex405) in response to control (0 MPa) or hyperosmotic stimulation (-0.75 MPa). All ratios are normalized to the mean value from t = 0 to 2 min. E, Intensity-based (Ex405 and Ex488) and ratio-based (Ratio) TIRF images of HyPer7-ROP6-expressing root epidermal cells in control (0 MPa) or after osmotic (-0.75 MPa) stimulation. F, Zoom in of a HyPer7-ROP6 nanodomain and its ratio (Ex488/Ex405) along the dashed line. G, Quantification of HyPer7-ROP6 oxidation level in control condition (0 MPa) or upon osmotic stimulation (-0.75 MPa) where signal in or out HyPer7-ROP6 nanodomains was segmented. H, Image segmentation based on lifetime values < or > than 2.9, of a cell expressing Hyper-7-ROP6 under osmotic stimulation. I, Quantification of HyPer7-ROP6 lifetime value in control condition (0 MPa) or upon osmotic stimulation (-0.75 MPa) among and outside osmotically-induced ROP6 nanodomains. Mean with error bars correspond to the 95% confident interval. For D * t-test, pvalue<0.05 at t=15 min. n is at least 5 seedlings from 3 independent replicates. For G and I, Letters show significant differences according to a one-way ANOVA and Tukey’s multiple comparisons test at p<0.05. For G, n cells is >53 from >9 seedlings from 3 independent replicates. For I, n cells is >9 from 6 seedlings from 2 independent replicates. For B, scale bar is 20 µm, for C scale bar is 50 µm, for E and H, scale bar is 10 µm.

Based on these results, we selected the HyPer7-ROP6 line for a detailed characterization of H_2_O_2_ signaling in response to osmotic stress. Five-day-old seedlings expressing HyPer7-ROP6 were transiently exposed to a hyperosmotic solution (-0.75 MPa) by perfusion. Since HyPer7 has an increased fluorescence excitation peak at 488lJnm when oxidized, and at 405 nm when reduced, we generated an Ex488/Ex405 ratio image to record its relative oxidation level. Within seconds of osmotic stimulus application, we observed a sharp increase in the Ex488/Ex405 ratio, followed by a plateau reached after 5 min (Figure 1C, D; Fig. S1 C). In contrast, washing out the hyperosmotic solution led to a decrease in the ratio, returning close to baseline levels (Figure 1C, D; Fig. S1 C). Perfusion with a control buffer resulted only in a slight but non-significant increase in the ratio (Figure 1C; Fig. S1 C). These findings indicate that osmotic stimulation leads to a significant and reversible increase in cytoplasmic oxidation. These findings show that osmotic stimulation leads to a pronounced and reversible increase in cytoplasmic oxidation revealed by the HyPer7-ROP6 probe. Taken together, our results demonstrate that HyPer7 is effective for detecting osmotic stress-induced alterations of cytosolic H_2_O_2_ levels, and that targeting of HPer7 to the PM as a ROP6 fusion does not compromise its oxidation sensitivity nor induce any observable phenotypic changes *in planta*.

### Osmotic stimulation induces oxidized nano environments at the PM

Since osmotic signaling induces the clustering and interaction of ROS-producing enzymes RBOHD and RBOHF with ROP6 (21), we hypothesized that this could lead to localized HyPer7 oxidation. Given that ROP6 nanodomains are clearly visible under Total Internal Reflection Fluorescence (TIRF) microscopy, we developed a method to quantify the degree of sensor oxidation under TIRF illumination (see Methods section). Under control condition, HyPer7-ROP6 displayed a uniform signal at the PM when excited at both 405 nm and 488 nm. We then generated ratio images between the sequentially imaged channels 488 nm/405 nm, and calculated a mean ratio value of 1.8 ± 0.24 (Figure 1E, Fig. S1 D and E). To validate our method, we applied increasing concentrations of exogenous hydrogen peroxide (H_2_O_2_), which resulted in a stepwise increase of the ratio values, consistent with dose-dependent oxidation of HyPer7-ROP6 (Fig. S1 D and E). These results demonstrate that ratiometric TIRF microscopy enables reliable monitoring of the oxidation dynamics of the HyPer7-ROP6 sensor at the PM of living plant cells.

After 15 minutes of osmotic stimulation, a portion of the HyPer7-ROP6 signal was found to accumulate in discrete clusters at the PM (Figure 1E). These clustering patterns closely mirror previous results obtained using GFP-ROP6 expressing lines (21, 28, 32), indicating that the HyPer7 fusion does not interfere with osmotic-induced ROP6 clustering. Interestingly, the ratio images revealed that the 488 nm/405 nm ratio values within these ROP6 nanodomains were significantly higher than those in adjacent membrane regions where HyPer7-ROP6 is also present (Figure 1E and F). To quantify this difference, we performed an image segmentation to separate fluorescence located in HyPer7-ROP6-enriched nanodomains from fluorescence outside these nanodomains. In osmotically treated cells, HyPer7-ROP6 ratio values were consistently elevated within nanodomains, while values outside these domains remained similar to those observed in untreated control cells (prior to any osmotic stimulation). This indicates that HyPer7 oxidation is spatially restricted and occurs specifically within nanodomains where ROP6 accumulates (Figure 1G).

To eliminate the possibility of changes in HyPer7 concentration affecting its photophysical properties and complicating our interpretation, we produced recombinant HyPer7 protein. Varying the protein concentration by more than twofold did not result in significant changes in the fluorescence ratio, confirming that the ratiometric signal from the sensor is not substantially influenced by its concentration (Fig. S1 F).

To further validate our findings, we measured HyPer7 fluorescence lifetime, which is less affected by sensor concentration than intensity. It remains responsive to oxidative changes as fluorescence lifetime increases in a dose dependent manner to exogenous application of H_2_O_2_ (Fig. S1 G). Minutes after osmotic stimulation, cells expressing HyPer7-ROP6 displayed an increased fluorescence lifetime, consistent with sensor oxidation (Fig. S1 H). Notably, pixel-wise lifetime segmentation revealed that elevated lifetimes were specifically confined to plasma membrane nanodomains (Figure 1H and I). These results indicate that osmotic changes induces a localized ROS accumulation restricted to ROP6-containing nanodomains, revealing the presence of spatially confined H_2_O_2_-rich nano environments at the cytoplasmic face of the PM.

To investigate whether this local H_2_O_2_ accumulation propagates deeper into the cytoplasm, we examined HyPer7 expressing lines under ratio-TIRF. Upon osmotic stimulation, no significant changes in mean HyPer7 ratio values nor distinct subcellular patterns were observed (Fig. S2 A and B). These observations suggest that the oxidized nano environments detected near the PM using HyPer7-ROP6 does not extend into the broader cytosolic space, supporting the existence of spatially restricted, H_2_O_2_ close to the cell surface. Given that our observations suggest H_2_O_2_ accumulation originates from discrete, nanodomains at the PM, we next investigated how this is modulated by the intensity of the osmotic stimulus. To this end, we applied a milder osmotic treatment (-0.26 MPa) to plants expressing HyPer7-ROP6. To quantify differences between osmotic strengths, we analyzed the distribution of pixel-wise ratio values specifically within HyPer7-ROP6-labeled nanodomains. Compared to the stronger stimulus (-0.75 MPa), the mild treatment resulted in a noticeable shift towards lower ratio values (Fig. S2 C-E). These data suggest a direct correlation between the osmotic stress intensity and nanodomains oxidation level.

To further examine the dynamic relationship between osmotic signal and localized oxidation, we performed washout experiments. 15 minutes after removal of the osmotic stimulus, HyPer7-ROP6 ratio values decrease significantly (Fig. S2 F-H). These trends became even more pronounced one hour after washout, indicating a full reversibility of nanodomain redox status in response to osmotic cues, even if ROP6 containing nanodomains are still present. Taken together, these results demonstrate that the strength of osmotic stimulation quantitatively correlates with the oxidation level below ROP6 nanodomains at the PM.

### The formation of oxidized nano-environments at the PM is signal-specific

We next asked whether the formation of osmotically induced oxidized nano-environments reflects a signal-specific cellular responses. To test this hypothesis, we examined the auxin signaling pathway, which also involves ROP6 clustering in both root and leaf epidermal cells (27, 31). Similar to GFP-ROP6, auxin treatment induced the formation of HyPer7-ROP6 nanodomains in root epidermal cells (Figure 2A). However, the HyPer7-ROP6 ratio values within these auxin-induced nanodomains did not differ significantly from those in surrounding membrane regions (Figure 2B). This finding was further confirmed at the tissue level, where auxin stimulation did not elicit any detectable changes in overall HyPer7-ROP6 oxidation (Figure 2C and D). These results suggest that the formation of oxidized nano-environments at the PM is specific to osmotic stress and does not arise from ROP6 clustering alone. Instead, it likely depends on the co-clustering of ROP6 with the ROS-producing enzymes RBOHD and RBOHF that only happens after osmotic stimulation and not auxin (21).

**Figure 2:**
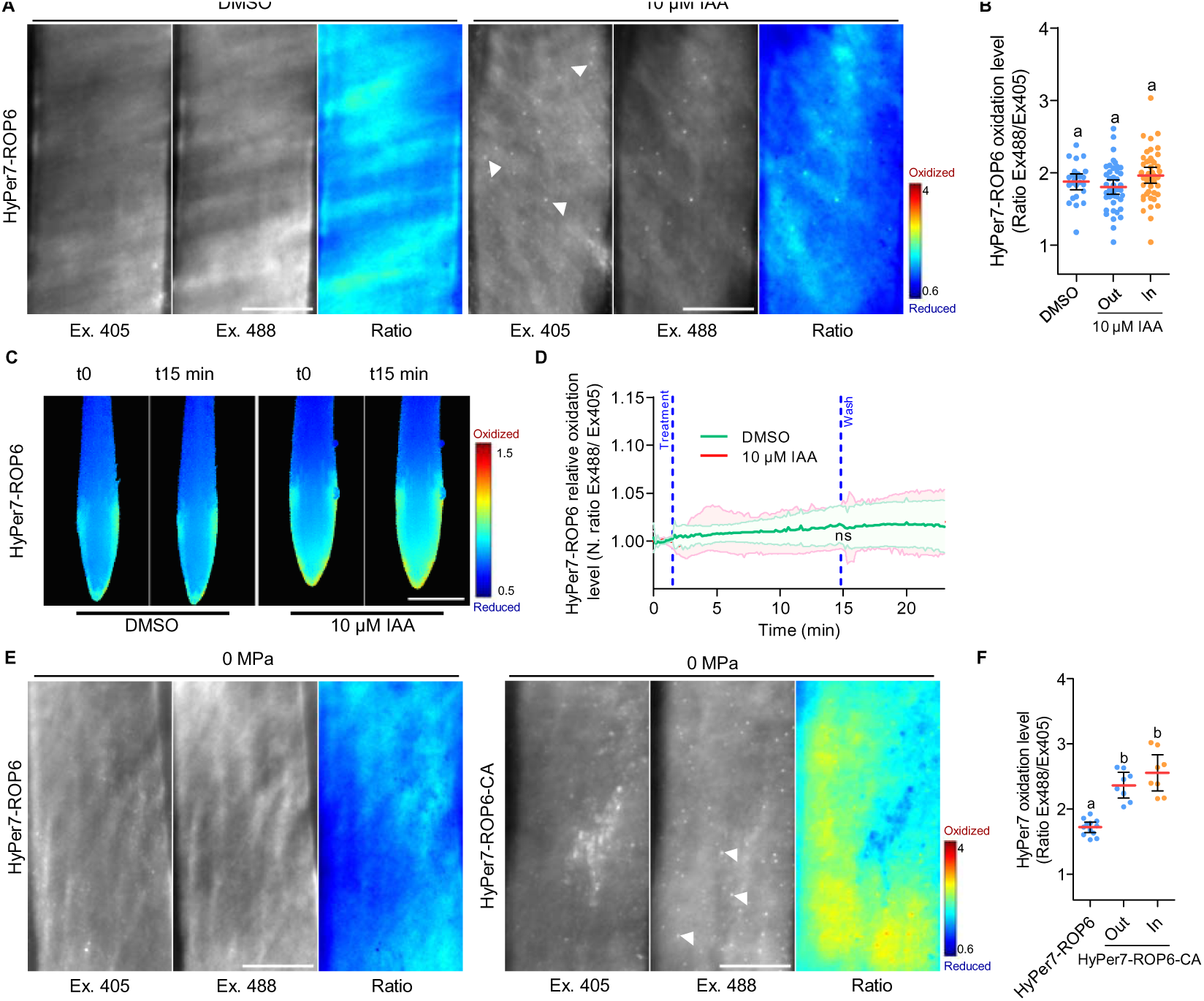
**Local HyPer7-ROP6 oxidation is not observed after auxin stimulation** A, Intensity-based (Ex405 and Ex488) or ratio-based (Ratio) TIRF images of HyPer7-ROP6–expressing root epidermal cells in control (DMSO) or after 10 µM auxin (IAA) stimulation. B, Comparison of HyPer7-ROP6 ratio values in control (DMSO) or after 10 µM auxin (IAA) stimulation in or out HyPer7-ROP6 nanodomains. C, Ratiometric images of 5 days old Arabidopsis plants expressing HyPer7-ROP6 in control condition or upon 10µM auxin stimulation. D, Typical time course showing HyPer7-ROP6 oxidation level (N. Ratio Ex488/Ex405) in response to control (DMSO) or after 10 µM auxin (IAA) stimulation. All ratios are normalized to the mean value from t = 0 to 2 min. E, Intensity-based (Ex405 and Ex488) and ratio-based (Ratio) TIRF images of HyPer7-ROP6 and HyPer7-ROP6-CA-expressing root epidermal cells in control condition. F, Comparison of HyPer7-ROP6-CA ratio values in control condition either in or out HyPer7-ROP6-CA nanodomains. Mean with error bars correspond to the 95% confident interval. For B and F letters show significant differences according to a one-way ANOVA and Tukey’s multiple comparisons test at p<0.05. For B, n >25 cells from >9 seedlings from 3 independent replicates. For F, n >8 cells from >4 seedlings from 2 independent replicates. For D, t-test, p-value=0.5058, n.s. means no significant differences n >5 seedlings from 3 independent replicates. A and E, Scale bar 10 µm. C, Scale bar 50 µm.

We then investigated to what extent ROP6 activation alone is sufficient to induce oxidized nano-environments at the PM. The expression of a constitutively active ROP6 variant carrying the G15V point mutation (ROP6-CA), which keeps the protein in its GTP-bound form, is known to promote ROP6 clustering and its interaction with RBOHD, even in the absence of external stimuli (21, 27, 31). As a result, ROP6-CA expression leads to basal H_2_O_2_ accumulation in plant cells (23–26, 33). Consistent with previous findings, HyPer7-ROP6-CA-expressing lines exhibited prominent nanodomains formation at the PM under non-stimulated conditions (Figure 2E). In this case, however, HyPer7-ROP6-CA ratio values were elevated both within and outside of nanodomains in ROP6-CA-expressing cells compared to control HyPer7-ROP6 lines (Figure 2F). These results indicate that constitutively active ROP6 is sufficient to induce both membrane clustering and HyPer7-ROP6-CA oxidation. However, the absence of spatially confined oxidation (i.e., similar ratio values inside and outside nanodomains) suggests that ROP6 activation alone does not recreate the condition in which the oxidized nano-environments are observed in response to osmotic stress. This implies that additional regulatory mechanisms are required to spatially restrict ROS production at the PM.

### H_2_O_2_ transport across the PM *via* aquaporin is necessary for establishing osmotically-induced oxidized nano-environments

Based on our results, we conclude that osmotic stimulation triggers the formation of oxidized nano-environments on the cytoplasmic side of the PM. However, it is well established that RBOH enzymatic activity consumes cytoplasmic NADPH to produce O_2_^-^in the extra-cellular space i.e. the apoplast. Given the short lifetime of O_2_^-^ (in the order of milliseconds), O_2_^-^ is generally assumed to be rapidly converted to H_2_O_2_ via dismutation and/or the action of superoxide dismutases (34, 35). Therefore, to explain the increased oxidation in the cytoplasm, we hypothesize the existence of a molecular mechanism that facilitates the translocation of an oxidizing agent, most likely H_2_O_2_, across the plasma membrane.

To test this hypothesis, we treated HyPer7-ROP6-expressing plants with catalase, a membrane-impermeable H_2_O_2_ scavenger, and then applied an osmotic stimulus. Catalase treatment did not abolish ROP6 nanodomain formation but markedly reduced HyPer7 oxidation in HyPer7-ROP6 nanodomains (Figure 3 A, C-D). These findings support a model in which RBOH-mediated O_2_^-^ production results in higher H_2_O_2_ levels in the apoplast, which must then cross the PM to oxidize HyPer7 in the cytoplasm. Conversely, co-application of exogenous H_2_O_2_ with osmotic stimulation enhanced oxidation levels of HyPer7-ROP6 nanodomains (Figure 3B, C-D). Interestingly, exogenous H_2_O_2_ in absence of any osmotic stimulation failed to generate oxidized nano-environment (Fig. S1 D). This suggests that osmotic signaling is required to enable or facilitate H_2_O_2_ diffusion through ROP6-containing nanodomains, reinforcing the idea that ROS signaling at the membrane is highly regulated and context-dependent.

**Figure 3:**
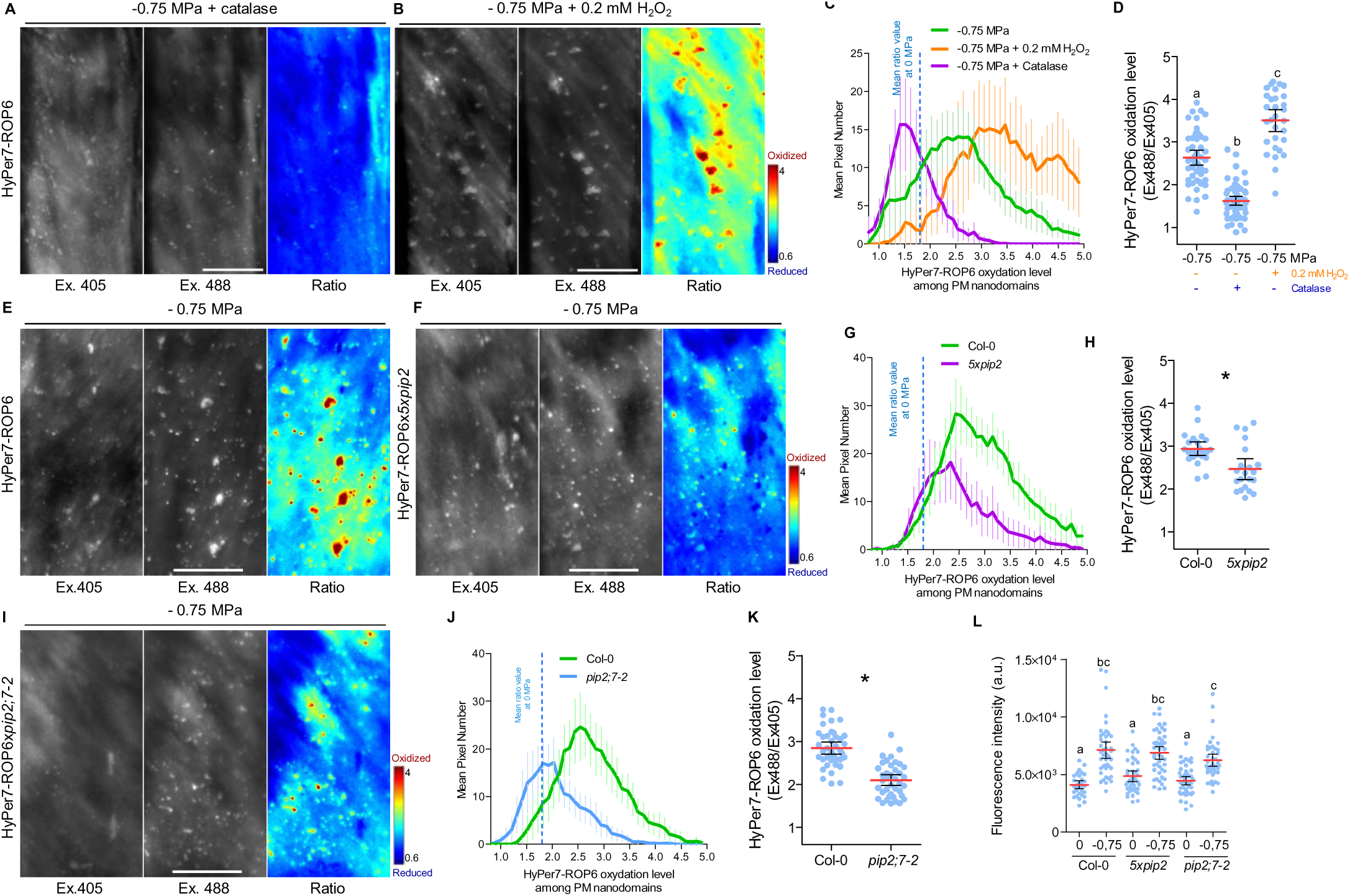
**Aquaporins are necessary for the establishment of osmotically-induced H_2_O_2_ nanoenvironments.** A-B, Intensity-based (Ex405 and Ex488) or ratio-based (Ratio) TIRF images of HyPer7-ROP6-expressing root epidermal cells upon strong osmotic stimulation (-0.75 MPa) co-treated with catalase, A or H_2_O_2_ (0.2 mM) B. C, Distribution of the pixel counts within HyPer7-ROP6 nanodomains as a function of their ratio values after osmotic stimulation (-0.75 MPa), and osmotic stimulation with catalase or exogenous H_2_O_2_ (0.2 mM). D, Corresponding quantification of the ratio values. E-F, intensity-based (Ex405 and Ex488) or ratio-based (Ratio) TIRF images of HyPer7-ROP6-expressing root epidermal cells upon osmotic stimulation (-0.75 MPa) in Col-0, E or in *5xpip2*, F. G, distribution of the pixel counts within HyPer7-ROP6 nanodomains as a function of their ratio values after osmotic stimulation (-0.75 MPa) in Col-0 or in *5xpip2*. H, Comparison of HyPer7-ROP6 ratio values after osmotic stimulation (-0.75 MPa) in Col-0 or in *5xpip2*. I, intensity-based (Ex405 and Ex488) or ratio-based (Ratio) TIRF images of HyPer7-ROP6-expressing root epidermal cells upon osmotic stimulation (-0.75 MPa) in *pip2;7-2.* J, distribution of the pixel counts within HyPer7-ROP6 nanodomains as a function of their ratio values after osmotic stimulation (-0.75 MPa) in Col0 or *pip2;7-2*. K, comparison of HyPer7-ROP6 ratio values after osmotic stimulation (-0.75 MPa) in Col0 or *pip2;7-2*. L, Dehydroethidium fluorescence quantification in col-0 or *5xpip2* in control condition or after 15 min treatment with -0.75 MPa buffer. Mean with error bars correspond to the 95% confident interval. For D and L, letters show significant differences according to a one-way ANOVA and Tukey’s multiple comparisons test at p<0.05. For D, n > 30 cells from >9 seedlings from 3 independent replicates. For L, n cells > 33 from 10 seedlings from 2 independent replicates. For H and K, t-test with a p-value=0,0013 and p-value=0,0001 respectively, n cells > 13 from >3 seedlings from 3 independent replicates. Scale bar is 10 µm.

To better understand the mechanism by which H_2_O_2_ might be diffusing from the apoplast to the cytoplasm within osmotically induced nanodomains, we explored the involvement of aquaporins. Indeed, aquaporins are membrane channel proteins primarily known for facilitating water transport but certain isoforms such as Arabidopsis PIP2;1; PIP2;2, PIP2;4, PIP2;5, PIP2;6, and PIP2;7 have also been shown to mediate the diffusion of H_2_O_2_ across membranes in various assays (36–38). The Plasma Membrane Intrinsic Protein 2 (PIP2) subfamily comprises a set of eight, highly homologous isoforms (39, 40), which have also overlapping functions. To overcome this, we generated a higher-order aquaporin mutant line from available single knockout lines, *5xpip2* (*pip2;1-1 pip2;2-3 pip2;4-1 pip2;6-3 pip2;7-2*), in which all the PIP2 isoforms that are highly expressed in the roots are knocked out (Fig. S3 A-D), (41, 42). We introduced by dipping HyPer7-ROP6 construct into this background to assess the impact of PIP2s into osmotic stress-triggered ROS signaling. First, we exposed Col-0 and *5xpip2*xHyPer7-ROP6 expressing plants to 10 mM H_2_O_2_ to fully oxidize the sensor. The apparent dynamic range of HyPer7-ROP6 for the ratio change between steady state and full oxidation after treatment of seedlings with 10 mM H_2_O_2_ was found similar in both backgrounds (Fig. S4 A), confirming that the sensor’s performances are not affected. HyPer7-ROP6-expressing roots in the *5xpip2* background exhibited a slower and reduced oxidation response following exposure to 100 µM exogenous H_2_O_2_ (Fig. S4 B). These results indicate that *5xpip2* mutants display reduced H_2_O_2_ permeability across the PM.

Under TIRF microscopy, HyPer7-ROP6 nanodomains were still able to form following osmotic stimulation but exhibited lower oxidation levels in the *5xpip2* background compared to Col-0 (Figure 3E-H). To rule out the possibility that this effect was due to reduced activity of the ROS-producing enzyme in *5xpip2*, we used dehydroethidium (DHE), a cell permeant dye that is primarily sensitive to O_2_^-^ *in vitro* and that labels the ROS including in the apoplast (43). Following osmotic treatment, both Col-0 and *5xpip2* roots showed comparable levels of DHE oxidation (Figure 3L), suggesting that RBOH activity remains intact in the *5xpip2* background. Together, these results demonstrate that both pharmacological (catalase) and genetic (*5xpip2*) disruptions of H_2_O_2_ diffusion from the apoplast to the cytoplasm significantly impact the oxidation status of osmotically-induced nano-environments at the PM. This supports the idea that the movement of H_2_O_2_ across the membrane is as a critical regulatory step in the localized signaling of ROS during osmotic stress.

To address whether the contribution of PIP2 aquaporins to osmotic signaling arises from the additive effects of multiple isoforms or is driven by specific individual PIP2 proteins, we generated a *pip2* triple mutant line from single knockout lines (3x*pip2*: *pip2;1-2/pip2;2-3/pip2;4-1*) expressing HyPer7-ROP6 by dipping (Fig. S3 B). Upon osmotic stimulation, *3xpip2*xHyper7-ROP6 show no difference in ROP6 nanodomain oxidation compared to control lines (Fig. S4 C-E). This result suggests that PIP2;1, PIP2;2 and PIP2;4 have a minor role in the formation of local ROS nano environments in root epidermal cells. PIP2;7 was previously shown to be abundant in root cells (41). In addition, mVenus-PIP2;7 expressed under the control of its endogenous promotor localizes at the PM of epidermal root tip cells like mCitrine-ROP6 does (44) (Fig. S4 F). Therefore, we generated a *pip2;7-2*xHyPer7-ROP6 line. We found that in response to an osmotic stimulation, whereas roots showed comparable levels of DHE oxidation than Col-0 (Figure 3L), HyPer7-ROP6 nanodomains were less oxidized, which closely resembled to the phenotype observed in the *5xpip2* mutant line (Figure 3 I-K and F-H). These results indicate that PIP2;7 is a major PIP2 isoform that facilitates H_2_O_2_ diffusion across the PM, leading to the formation of H_2_O_2_ nano-environments in the cell epidermis. Thus, the *pip2;7* mutant provides a unique opportunity to test the functional importance for osmotic signaling of the localized H_2_O_2_ nano-environments on the cytosolic side of the PM.

At the cellular level, osmotic signal trigger pronounced anisotropic cell expansion (45, 46) a response partially regulated by ROP6 (21). Consistent with this, we found that the double mutant of ROS-producing enzymes, *rbohd/f*, displays an impaired cell swelling response similar to that observed in *rop6-2* plants (Figure 4A, B). We next tested this response in aquaporin mutants; in which the ROS levels are increased after osmotic stress in the apoplast but not in the cell cortex. Both the *5xpip2* and *pip2;7-2* mutants showed reduced cell swelling in response to osmotic stimuli than the wild type control. By contrast, the *3xpip2* mutant, which did not impact the formation of oxidized nanodomains, showed a cell swelling similar the wild type after osmotic stress, suggesting that the H_2_O_2_-oxidized nano-environment positively regulates anisotropic cell growth in response to osmotic stimulation (Figure 4A, B).

**Figure 4:**
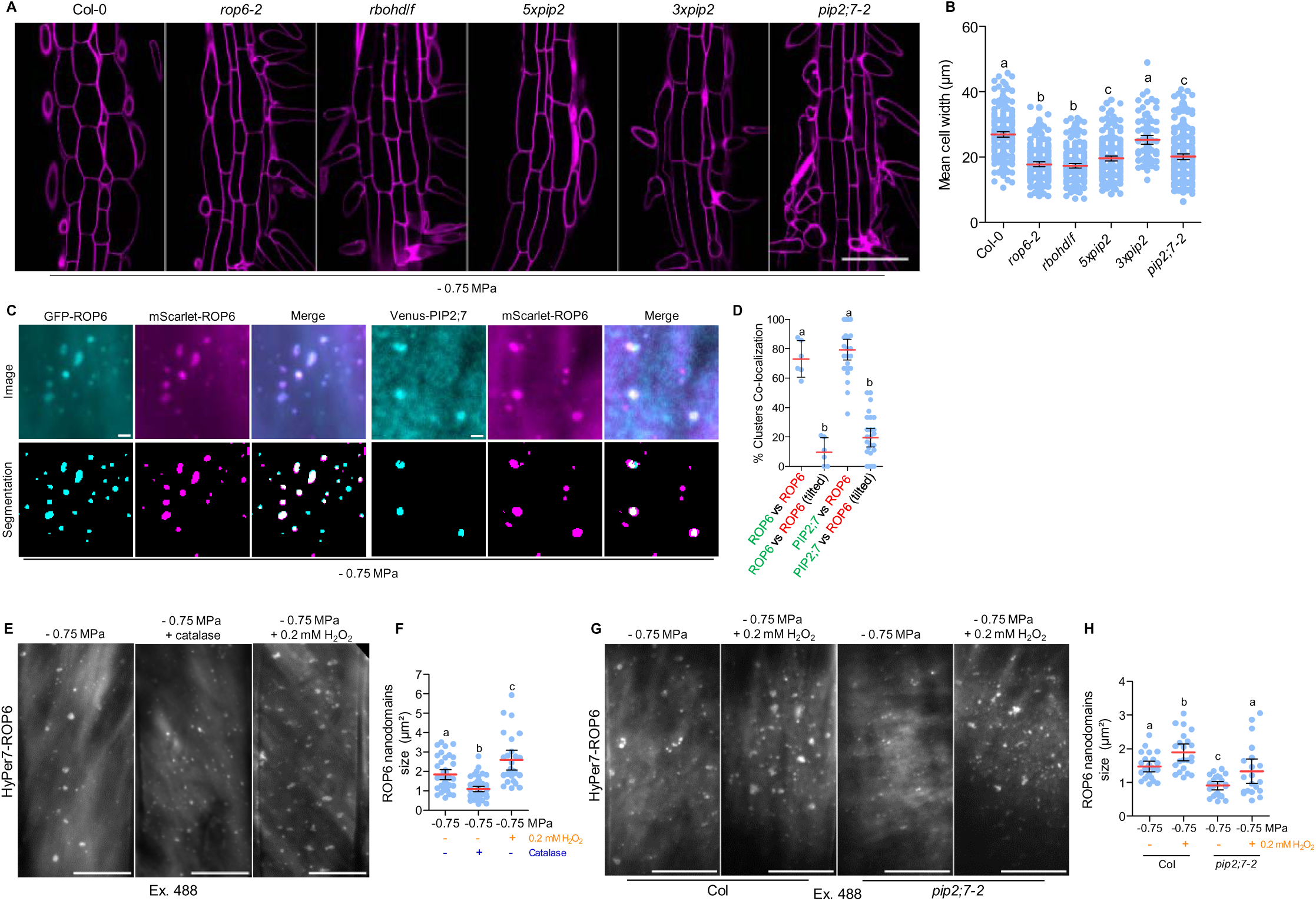
**PIP2;7 colocalizes in osmotically-induced ROP6 nanodomains, plays a role in ROP6 clustering and participate in anisotropic growth** A, Confocal images of root epidermal cells of Col-0, *rop6-2*, *rbohd/f*, *5xpip2*, *3xpip2* and *pip2-7* transferred for 24h at -0.75 MPa on vertical petri dishes. Quantification of cell width is shown in B. C, Dual-TIRF images of root epidermal cells expressing p35S:GFP-ROP6 x pPIN2:mScarlet-ROP6 or pPIP2;7:mVenus-PIP2;7 x pPIN2:mScarlet-ROP6 upon osmotic stimulation. Image segmentation (shown below) was performed to select exclusively pixels among clusters, used to calculate the percentage of colocalization shown in D. As control, 90° tilt was done on one of the two channels. E, TIRF images of root epidermal cells expressing HyPer7-ROP6 (Ex 488) upon osmotic stimulation and co-treated with catalase or 0.2 mM exogenous H_2_O_2_. ROP6 nanodomains size was quantified after image segmentation and is shown in F. G, TIRF images of root epidermal cells expressing HyPer7-ROP6 (Ex 488) upon osmotic stimulation co-treated or not with 0.2 mM exogenous H_2_O_2_ in Col-0 or *pip2;7* background. ROP6 nanodomains size was quantified after image segmentation and is shown in H. Mean with error bars correspond to the 95% confident interval. Letters show significant differences according to a one-way ANOVA and Tukey’s multiple comparisons test at p<0.05. In A, n > 88 cells from > 12 seedlings from 2 independent replicates. In D, n cells >5 from > 4 seedlings from 2 independent replicates. In F, n cells >26 from >9 seedlings from 3 independent replicates. In H, n cells >20 from >9 seedlings from 3 independent replicates. For A, scale bar is 50 µm, For C, scale bar is 1 µm. For E and G scale bar is 10 µm.

### PIP2;7 co-localizes in ROP6-nanodomains and the resulting sub-membrane oxidation reinforcing ROP6 clustering

Thus, our data suggest a functional link between ROP6 and PIP2;7 in establishing osmotically-induced oxidized nano-environments. Thus, we investigated by dual-color TIRF microscopy whether PIP2;7 upon osmotic stimulation assembles in nanodomains and whether it co-localizes with ROP6. Minutes after osmotic treatment, both mVenus-PIP2;7 and mScarlet-ROP6 formed nanodomains that showed significant co-localization (Figure 4 C, D, Fig. S4 G). These observations suggest that ROP6 nanodomains, in addition to RBOHD/F, include PIP2;7, likely forming a functional complex that coordinates H_2_O_2_ production and transport at specific sites of the PM.

Pharmacological and genetic perturbation of H_2_O_2_ often lead to a change in ROP6 nanodomains shape as observed on TIRF experiments. Specifically, exogenous application of H_2_O_2_ combined with osmotic stimulation results in larger ROP6 nanodomains (Figure 4E, F). Conversely, limiting apoplastic H_2_O_2_ either through catalase treatment or by impairing its transport in the *pip2;7-2* mutant, leads to smaller ROP6 nanodomains (Figure 4F, H). To test whether H_2_O_2_ accumulation is a limiting factor for ROP6 nanodomains size, we treated the *pip2;7-2*xHyPer7-ROP6 line with exogenous H_2_O_2_, which partially restored the nanodomains size (Figure 4G, H). These results suggest that H_2_O_2_-enriched nano-environments may contribute directly to a feedforward regulatory loop that controls the size of ROP6 nanodomains.

## DISCUSSION

The presented work demonstrates that ROP6 is activated by osmotic stress, leading to the formation of ROP6-containing nanodomains at the PM. These nanodomains are associated with discrete oxidized nano-environments at the cytosolic side of the PM. Such process requires diffusion of H_2_O_2_ across the membrane, in a PIP2;7-aquaporin dependent manner. In turn, the localized oxidation just below the PM feeds back to regulate the formation of ROP6-containing nanodomains, partially controlling the plant response to osmotic stimulation, as cell isotropic growth. In addition, this model exemplifies that protein clustering in nanodomains could lead to the formation of a local gradients of small diffusible molecules like H_2_O_2_ that is controlled both across and in the plane of the membrane.

Our observations indicate that in addition to the organization of ROP6 and RBOH in nanodomains, aquaporin-dependent H_2_O_2_ diffusion across the PM also appears to play a critical role. Interestingly, aquaporins, which allow H_2_O_2_ diffusion, both in heterologous systems and *in planta*, are regulated by kinases similar to those that modulate RBOH activity. For example, both RBOHF and PIP2;1 are phosphorylated by the kinase OST1/SnRK2.6 (37, 47, 48). The formation of RBOH/PIP nanodomains co-regulated by SnRK2 for a coordination between RBOH-dependent ROS production and H_2_O_2_ diffusion across the membrane regulated by PIPs is an appealing hypothesis, which would need further analyses.

In previous studies, cerium chloride staining combined with electron microscopy was elegantly used to visualize H_2_O_2_ accumulation in plant cells. For example, cryptogein, a *Phytophthora*-derived elicitor, induces RBOH-dependent H_2_O_2_ accumulation in discrete foci at the PM when applied to tobacco cells (49). Similar observations were made in the context of lateral root emergence: a discontinuous pattern of H_2_O_2_ accumulation were observed at the endodermal PM (50). For those two examples, the exact localization of the H_2_O_2_ accumulation: facing the cytoplasm or the apoplast remains to be determined. More recently, a highly defined localization of H_2_O_2_ accumulation in the apoplast was described during the lignification of the Casparian strip (51, 52). These observations suggests that localized H_2_O_2_ accumulation at the PM extends way beyond the specific context of osmotic stress responses and may occur in diverse cellular processes. Although it remains to be demonstrated if this localized H_2_O_2_ accumulation results in spatially confined changes in the cytoplasmic redox environment. If yes, this could contribute to the activation of specific downstream signaling cascades mediated by H_2_O_2_.

Provided protein thiols are sufficiently reactive, H_2_O_2_ might oxidize target proteins directly. More likely, however, decoding hubs consisting of highly sensitive H_2_O_2_-sensing proteins capable of transmitting the primary oxidation to protein thiols and the respective downstream targets may exist. In this case, protein-protein interactions and proximity would define the specificity of such oxidation reactions. Detoxification of H_2_O_2_ in the ascorbate-glutathione pathway has been proposed to trigger changes in the local glutathione redox potential (*E*_GSH_), which may cause changes in glutathionylation of target proteins in a reaction catalyzed by glutaredoxins (GRXs) (53). Alternatively, peroxidases acting on H_2_O_2_ may also function as protein-thiol oxidases as it has recently been shown for the role of cytosolic peroxiredoxin PRXIIB in transmitting the primary oxidation induced by pathogens to the PP2C protein phosphatases ABI1/2 (54). Disulfide formation on ABI1/2 inhibits the phosphatases and ultimately leads to stomatal closure (54). Similarly, it was shown earlier that the glutathione peroxidase Gpx3 (also known as Orp1) in *Saccharomyces cerevisiae* can transmit a H_2_O_2_ signal to the transcription factor Yap1 to switch it to the active oxidized form (55). The efficiency and specificity of such transmission systems rely on protein proximity and it can thus be postulated that appropriate scaffold proteins or other structural means exist to mediate proximity between the peroxidase as the H_2_O_2_ sensor and target proteins. Therefore, we can speculate that localization of GRX, TRX and potentially GPX in specific nanodomains would determine signal transduction from the membrane.

Recently, in yeast, H_2_O_2_ concentration changes were carefully mapped, revealing dynamic and highly specific variations in H_2_O_2_ availability at the level of individual proteins and protein complexes (56). Our data closely mirror this phenomenon, supporting the idea that ROS signaling is spatially discretized within cells. In line with this, we propose that nanodomains composed by RBOHs, ROPs, and PIPs represents a fundamental unit for PM–localized ROS signaling in plants.

## MATERIALS AND METHODS

### Plant material and growth condition

*Arabidposis thaliana* ecotype (Col-0) was used as wild-type control. The following lines were previously published: pPIP2;7:mVenus-PIP2;7 (44), p35S:RFP-ROP6 (21) and *pipi2;7-2* (42). The multiple *pip* mutants *5xpip2* and *3xpip2* were obtained by genetic crossing of *pip2;1-2* (SM_3_35928; (57)), *pip2;2-3* (SAIL_169A03; (58)), *pip2;4-1* (SM_3_20853), *pip2;6-3* (SALK_092140; (59)), and *pip2;7;2*; all insertion lines were obtained from the Arabidopsis stock centers (60). ROP6 (21), *5xpip2*, *3xpip2* and *pipi2;7-2* (42). The lines p35S:HyPer7-ROP6, p35S:HyPer7-ROP6-CA, *5pip2*xp35S:HyPer7-ROP6 and *3pip2*xp35S:HyPer7-ROP6 were generated by floral dipping. Crosses *pip2;7-2*xp35S:HyPer7-ROP6, p35S:RFP-ROP6xpUBQ:mScarlet-ROP6 and pPIP2;7:mVenus xpUBQ:mScarlet-ROP6 were done in this study. Seeds were surface-sterilized and sow on squared petri dishes containing 50 mL of half-strength Murashige and Skoog medium (MS/2) supplemented with 1% (w/v) sucrose, 2.5 mM MES-KOH (pH 6) and 0.8% agar type E. Seedlings were grown vertically for five days under a 16-h light/8-h dark cycle, 70% relative humidity, and a light intensity of 200 µmol·m⁻²·s⁻¹.

### Cloning

HyPer7 clones were generated by Golden Braid (61). In brief, the coding sequence of HyPer7 (30) and ROP6 (At4g35020) were PCR amplified and cloned in pUPD2 vectors. The G15V mutation in ROP6 sequence were introduced by PCR mutagenesis to generate pUPD2-ROP6-CA. Binary vector were made from the assembly of the following parts pUPD2:p35S, pUPD2-HyPer7, pUPD2-ROP6, pUPD2-ROP6-CA and pUPD2:tNos to create pDGB3a1 p35s:HyPer7:tNos, p35s:HyPer7-ROP6:tNos and p35s:HyPer7-ROP6-CA:tNos, to generate transcriptional units. To generate pPIN2:mScarlet3-ROP6 a modified GreenGate cloning system was employed (62). Briefly, GreenGate entry vectors were modified by replacing the *caR-ccdB-lacZa* cassette with *lacZa* flanked by SapI restriction sites that overlap with the existing BsaI restriction sites. The shuttle vectors were used in a combined restriction-ligation reaction to clone DNA fragments into the shuttles and create the six modules required for assembly of the binary vector. The following DNA fragments were cloned into their respective shuttle vector: promoter of the *PIN2* gene (module A), mScarlet3 fluorophore (63); module B), *ROP6* genomic sequence (module C), 3’-UTR of *ROP6* (module D). A combined terminator of the HSP and rbcS E9 terminator sequences was cloned into the classical GreenGate entry vector pGGE000 (module E) (see table for primer sequences). For the binary vector, the backbone of pDGE651 (64) was combined with the orange selection of the GoldenGate vector pAGM4673. All clones were Sanger sequenced. See Fig S5 for primer sequence.

### Osmotic and pharmacological treatments

Osmotic stimulation was applied by first incubating seedlings for 15 min in liquid resting buffer (MS/2, 2.5 mM MES-KOH, pH 6 and 1% w/v sucrose). Seedlings were then transferred to sorbitol buffer composed by the resting buffer supplemented with 150 or 300 mM sorbitol to impose osmotic shocks of -0.26 or -0.75 MPa, respectively. 40u/ml catalase was added to both resting (15 min) and sorbitol (15 min) buffers, whereas 0.02 mM, 0.1, 0.2 or 1 mM H_2_O_2_ was added only for 15 min either to the resting buffer or to resting buffer supplemented with 300mM sorbitol. Auxin treatments were done for 20 min at 10 µM diluted from a stock at 100 mM raised in DMSO.

### HyPer7 sensor oxidation range and dynamics in a multiwell plate reader

Experiments were performed as described in (65). Briefly, pools of 10-day-old Wild-type or *5xpip2* seedlings expressing either HyPer7-ROP6 or HyPer7-Cyto were placed in independent wells of 96-well plate pre-loaded with 180 µL of imaging buffer. The redox state of the sensors was followed as the fluorescence ratio between two channels collected sequentially (Channel 1: Ex = 400 ± 5 nm / Em = 520 ± 5 nm; Channel 2: Ex = 482 ± 8 nm / Em = 520 ± 5 nm), using a CLARIOstar plate reader (BMG Labtech, Ortenberg, Germany).

### Ratiometric microscopy

To monitor HyPer7 oxidation dynamics at the root tissue level over time, seedlings were mounted in a perfusion chamber coupled to a ratiometric imaging setup. The setup consisted of a Zeiss Axiovert 200M microscope equipped with a ORCA-Fusion BT (Hamamatsu) camera, a lambda 10B wheel (Sutter Instrument) and LEDHub (Omicron) illumination source, all controlled via the electronic control unit and software from Inscoper. Images were acquired using a 20x/0.5 objective, with excitation set to 385 nm (370-415) or to 470 (430-540) and emission collected at 535/30 nm. Sequential images for both channels were acquired every 10 seconds, and 470/385 ratio images were generated to quantify HyPer7 oxidation.

Prior to treatment, samples were acclimated in resting buffer within the incubation chamber for 30 min to allow the plant to rest and minimize light-induced ROS production. Treatments were applied by complete buffer replacement through three successive perfusions. Washes were performed using the same method.

### Image processing for ratiometric microscopy

A custom ImageJ macro was used to process two-channel fluorescence image stacks and compute ratiometric profiles. Each multichannel image was split into separate 405 nm and 488 nm stacks, and background-corrected using a manually defined ROI. Images were converted to 32-bit format, and low-intensity pixels (below 300 for 405 nm and 200 for 488 nm) were set to NaN to exclude background noise. The resulting stacks were saved together with their Z-axis intensity profiles. A ratio image (488/405) was then generated and exported along with its Z-axis profile for quantitative analysis.

### Ratiometric total internal reflection fluorescence microscopy

A custom TIRF system was configured on an inverted Zeiss microscope equipped with a 100x/1.45 oil-immersion objective. An additional telescope placed in the imaging path provides a final magnification of 150x at the camera sensor. The TIRF angle was manually adjusted to give the best signal to noise ratio. Sequential acquisitions were performed using laser excitation at 488 nm and 405 nm with images collected at 525/25 nm. Image series were collected at three z-planes (0.125 µm step). For each plane, 20 frames per channel were acquired sequentially. To correct for field inhomogeneities, which are inherent to TIRF illumination and vary with wavelength and incidence angle, flat-field correction images with 200 frames per channel were acquired.

### Image processing for ratiometric TIRF

A custom ImageJ macro was used to batch process multi-channel fluorescence images for ratiometric analysis (Fig S6). Each image file was split into two channels. For each channel, a Z-projection based on average intensity was generated and converted to 32-bit format. To correct for uneven illumination, each projection was divided by the corresponding flat-field correction image. The corrected 488 nm and 405 nm projections were then divided to generate a pixel-by-pixel ratio image (488/405), which was displayed using the “Jet” LUT and a standardized brightness/contrast range (0.6–4.0). The resulting ratio images, and the separated 405 nm and 488 nm projection images were then saved in designated output folders.

For quantification of clustering or ratio values, a crop of 100 pixels square was realized on each image. Image segmentation to discriminate pixels among nanodomains was realized using machine learning assisted method (with ilastik software, (66)). Created segmented images where then used to determine ROIs corresponding to pixels among or outside nanodomains.

To determine the distribution of ratio values of the pixels in HyPer7-ROP6 nanodomains, a custom ImageJ macro was used to create a binary mask on segmented images in order to created composite ROI containing all those pixels. An intensity histogram of 41 bins, range 0.8–5 (range covering the ratio values observed with HyPer7-ROP6 upon osmotic stimulation) was then generated for each ratio image. The different macros used for image processing are provided in the supplement (Supp Macros File).

### In vitro HyPer7 characterization

HyPer7 was produced *in vitro* and purified using His-tag according to Ugalde et al., 2021. Purified sensor samples were imaged on the same TIRF setup, using both 488 nm and 405 nm as excitation and 525/25 for emission. To estimate sensor concentration and oxidation state in serial dilution, we use total intensity and 488/405 ratio, respectively. Three different sensor dilutions were imaged to achieve fluorescence signals comparable to those observed inside and outside osmotically induced HyPer7-ROP6 PM nanodomains in living samples.

### Lifetime determination of HyPer7-ROP6 *in planta*

Five-day-old Arabidopsis seedlings expressing HyPer7-ROP6 treated or not with -0.75 MPa solution were imaged on a Leica Stellaris 8 Falcon confocal microscope coupled with an 80 MHz WLL laser tuned at 488 nm. Images of epidermal cells at the root elongation zone were collected using a 40x/1.1 water immersion objective. The fluorescence was collected between 500 and 550 nm. The lifetime of HyPer7-ROP6 sensor was estimate by phasor plot with a wavelet filter and pixels with a total photon count bellow five have been remove of the analysis.

### Confocal laser scanning microscopy

Localization study of p35S:HyPer7 (Cyt), p35S:HyPer7-ROP6, pROP6:mCit-ROP6 and pPIP2;7:mVenus-PIP2;7 was monitored in five-day-old Arabidopsis seedlings stably expressing the respective constructs. Epidermal cells in the root elongation zone were imaged using a 40x1.1 water immersion objective on a Leica SP8 confocal microscope. Excitation was set to 488 nm, and emission was collected at 525/22 nm. Seedlings were grown on vertical Petri dishes containing standard MS/2 medium as described above and acclimated in liquid resting buffer for 15 min prior to imaging.

### Dual color total internal reflection fluorescence microscopy

Dual-color TIRF imaging was performed using the same custom-made TIRF setup described previously. For GFP or mCitrine imaging, excitation was provided by a 488 nm laser (OBIS LX 488-50, Coherent Inc.; 40 mW), and emission was collected using a 525/ 22 nm filter. For mScarlet imaging, excitation was provided by a 561 nm laser (SAFIR 100 mW), and emission was collected using a 600/20 nm filter. To avoid bleed-through of fluorescence signal between GFP/Venus & mScarlet, single-color plant lines were imaged in each condition, confirming that each fluorophore was detected in well separated emission channels without detection of any bleed-through signal. Image segmentation was performed as describe for Ratiometric TIRF imaging to identify protein clusters, and merged to quantify the percentage of colocalizing particles. As a negative control, the red channel was tilted by 90°, the images were re-merged, and the resulting percentage of colocalization was determined.

### Western blotting

Root tissue from 12-day-old Col-0, *5xpip2*, *3xpip2* and *pip2;7-2* plantlets were ground with liquid nitrogen to a fine powder and resuspended in 100 mg/ml of extraction buffer (250 mM Tris-HCl pH6.8, 5% SDS, 40% Glycerol, 0.02% bromophenol blue and 250 mM DTT). Western blot analysis was performed with antibodies diluted at 1/5,000 for anti-PIP2 and anti-PIP2;7 (Agrisera AS22 4812); and 1/20,000 for HRP conjugated anti-rabbit in blocking buffer (2% not fat milk, 0.02% Tween 20, 1x TBS).

### Cell swelling assay

We performed the cell-swelling assay on 7 days old Arabidopsis plants. 6 days grown seedlings (MS/2, 2.5mM MES-KOH pH5.8, 1% sucrose, 0.8% Agar Type E) were transferred for 24h on the same media supplemented with 300 mM Sorbitol. Plants were stained with propidium iodide (1 µl/ml) for 3 min in liquid buffer (MS/2, 2.5mM MES-KOH pH 5.8, 1% sucrose) supplemented with 300 mM Sorbitol. After a 2 min wash, the plants were imaged using a confocal microscope.

### Statistics

In each experiment, 8 to 10 cells were studied from five to seven different seedlings. All experiments were independently repeated two to three times. Each cell was analyzed individually to include the maximum of the biological variation. Data are expressed as mean ± 95% confidence interval. Analysis of variance (ANOVA) followed by Tukey test was done, and letters indicate significant differences among means (*P* < 0.05). Statistical analyses such as ANOVA followed by a Tukey post hoc test and *t* test were done in GraphPad Prism.

## ACKNOWLEDGMENTS

We thank the Montpellier Ressources Imagerie (MRI) and the Histocytology and Plant Cell Imaging Platform for providing the microscope facility (PHIV). We acknowledge the MARS imaging facility, member of the national infrastructure France-BioImaging (FBI) supported by the French National Research Agency (ANR-10-INBS-04). We thank the Root Phenotyping Platform (PhR) for their support with non-destructive root imaging and kinetic analysis of root system architecture. This work was supported by the Agence National de la Recherche (ANR) through the project CellOsmo (ANR-19-CE20-0008) to AM and joint funding by ANR and the Deutsche Forschungsgemeinschaft (DFG) through the project Nano-ROS (ANR-24-CE92-0024 to AM and DFG 545516367 to AJM). Work in YJs laboratory is supported by the European Research Council (ERC) under the European Union’s Horizon 2020 research and innovation program (Grant Agreement No 101001097-LIPIDEV) to YJ, by an EMBO long-term fellowship (ALTF 750-2023) and Marie Curie Sklodowska Action (HORIZON-TMA-MSCA-PF-EF No 101151872-SONA) to PS. KJ is grateful for a doctoral fellowship awarded by the German Academic Exchange service (DAAD, Germany).

