## Supplemental figures for "Nanoscale regulation of ROS signaling at the plasma membrane tunes the plant response to osmotic stress"

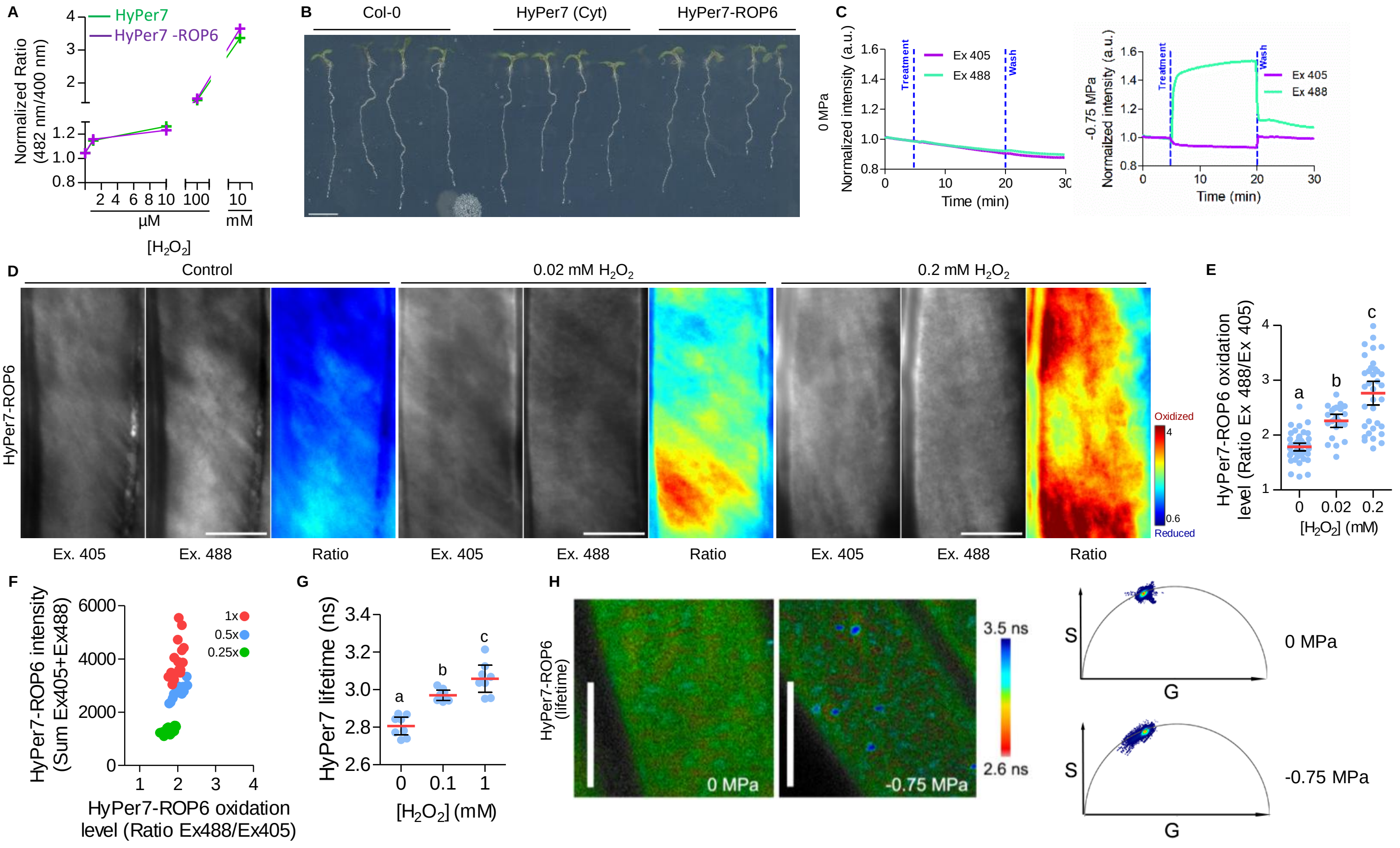


**Figure S1: ROP6 fusion does not affect HyPer7 sensitivity to $H_2O_2$ nor plant development**

A, HyPer7 (green) and HyPer7-ROP6 (purple) ratio in control condition after progressive oxidation with 1, 10, 100µM and 10 mM exogenous $H_2O_2$. B, 5 days old Col-0 and transgenic lines expressing HyPer7 and HyPer7-ROP6, grown on MS/2 vertical petri dishes. C, Typical time course showing normalized intensity of 488 and 405 channels in 5 days old plantlets expressing HyPer7-ROP6 upon control condition (0 MPa) or upon osmotic stimulation (-0.75 MPa). D, Intensity-based (Ex405 and Ex488) and ratio-based (Ratio) TIRF images of HyPer7-ROP6-expressing root epidermal cells after application of increasing concentration of exogenous $H_2O_2$ and its respective quantification in E. F, Ratiometric (488/405) and total intensity (405+488) quantification of *in vitro* HyPer7 at various concentrations (1x, 0.5x, and 0.25x). G, HyPer7-ROP6 lifetime quantification of HyPer7-ROP6 expressing root epidermal cells after application of different $H_2O_2$ concentrations. H, Confocal micrographs showing HyPer7-ROP6 lifetime in control condition (0 MPa) or upon osmotic stimulation (-0.75 MPa) and their corresponding phasor plots.

*Mean with error bars correspond to the 95% confident interval. Letters show significant differences according to a one-way ANOVA and Tukey's multiple comparisons test at $p<0.05$. In E, n cells > 24 from >9 seedlings from 3 independent replicates. In G, n cells > 9 from >6 seedlings from 2 independent replicates For D and H, scale bar is 10 µm.*

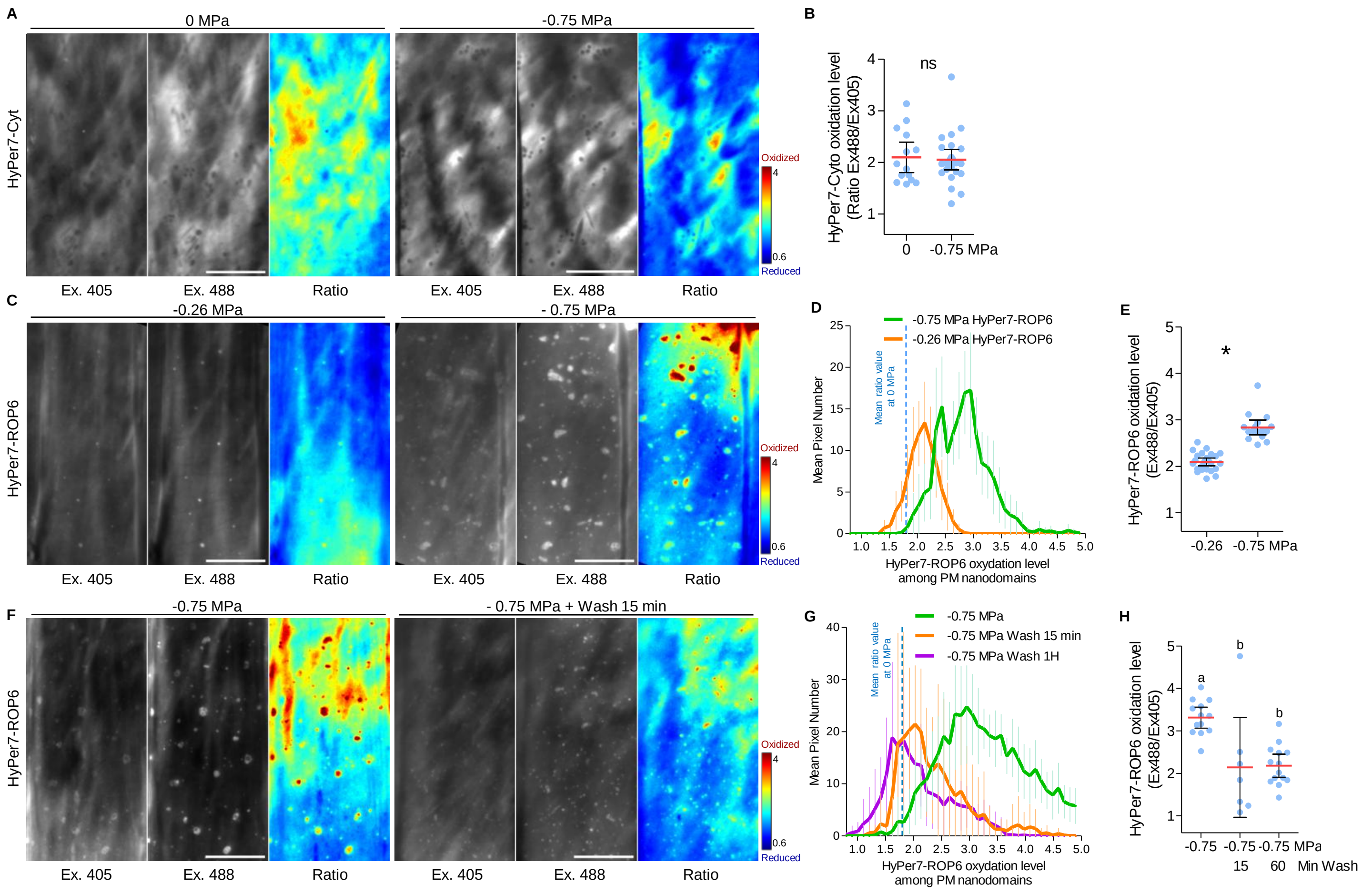


**Figure S2: HyPer7-ROP6 localized oxidation is dependent on the strength of the osmotic shock and is not observed with the soluble HyPer7**

A, Intensity-based (Ex405 and Ex488) or ratio-based (Ratio) TIRF images of HyPer7 expressing root epidermal cells in control condition (0 MPa) or after an osmotic stimulation (-0.75MPa) and its respective quantification in B. C, Intensity-based (Ex405 and Ex488) and ratio-based (Ratio) TIRF images of HyPer7-ROP6 expressing root epidermal cells after a mild (-0.26 MPa) or stronger (-0.75 MPa) osmotic stimulation. D, Distribution of the pixel counts within HyPer7-ROP6 nanodomains as a function of their ratio values after a mild (-0.26 MPa) or stronger (-0.75 MPa) osmotic stimulation. E, Comparison of HyPer7-ROP6 ratio values after a mild (-0.26 MPa) or stronger (-0.75 MPa) osmotic stimulation. F, Intensity-based (Ex405 and Ex488) and ratio-based (Ratio) TIRF images of HyPer7-ROP6 expressing root epidermal cells after osmotic stimulation (-0.75 MPa) or after a 15min wash with control buffer. G, Distribution of the pixel counts within HyPer7 nanodomains as a function of their ratio values after osmotic stimulation (-0.75 MPa) or after a 15min and 1h wash with control buffer. H, Comparison of HyPer7-ROP6 ratio values after osmotic stimulation (-0.75 MPa) or after a 15 min an 1H wash with control buffer.

*Mean with error bars correspond to the 95% confident interval. For B t-test, p-value=0,7692, n.s. means no significant differences n cells >14 from >3 seedlings from 3 independent replicates. For E t-test, p-value<0,0001 n cells >16 from >3 seedlings from 3 independent replicates. For H, letters show significant differences according to a one-way ANOVA and Tukey's multiple comparisons test at p<0.05. n cells > 7 from >3 seedlings from 2 independent replicates. Scale bar is 10 µm.*

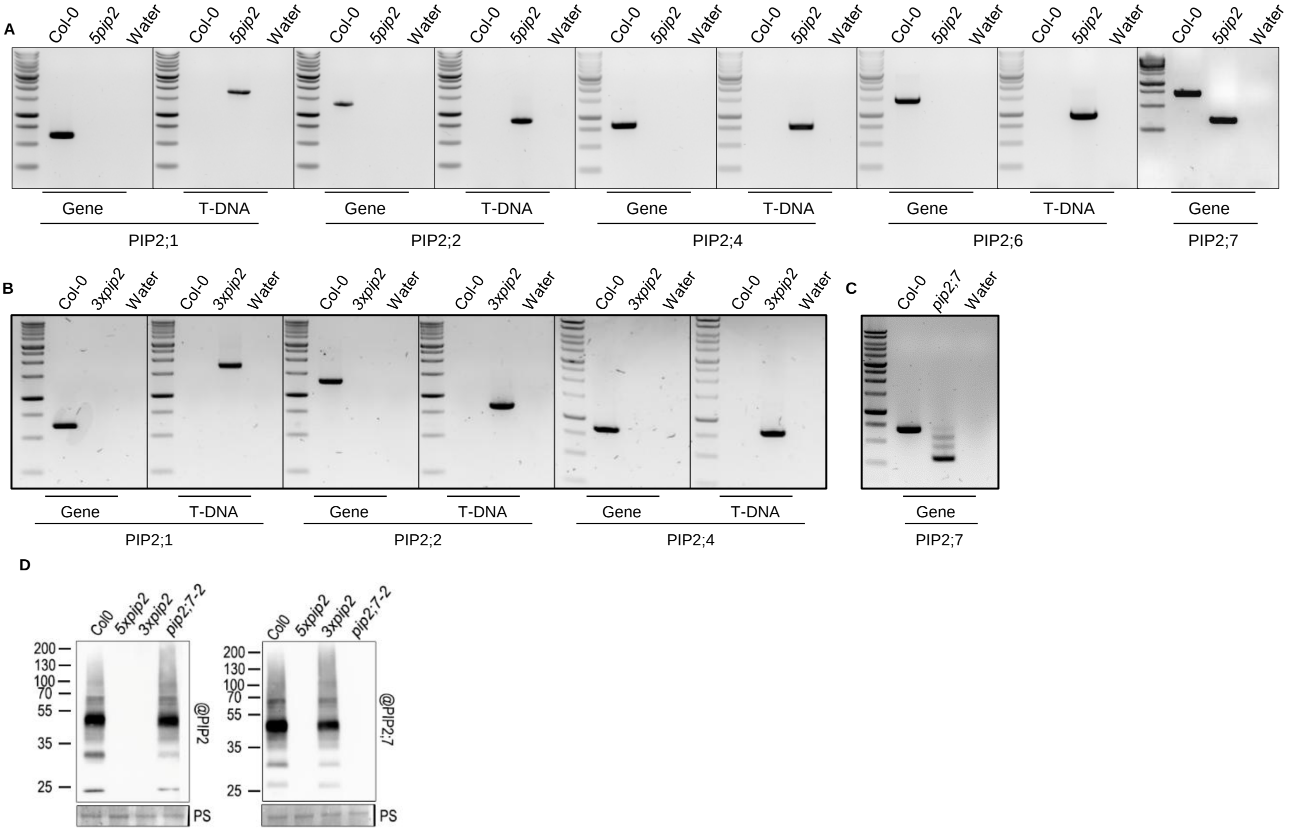


### Figure S3: Genetic characterization of PIP2 mutants

A, PCR electrophoresis for wild-type genes (*PIP2;1*, *PIP2;2*, *PIP2;4*, *PIP2;6*, *PIP2;7*) and corresponding T-DNA insertions or CrispR mutations in Col-0 and the quintuple PIP2 mutant (*5xpip2*). Same is shown for the triple mutant PIP2 (*3xpip2*) in B, and for the simple CrispR mutant *pip2;7-2* in C. D, Western blot from root protein extract from Col0, *pip2;7-2*, *3xpip2* or *5xpip2* and revealed with antibody against PIP2 that recognizes PIP2;1, PIP2;2 and PIP2;3 or against PIP2;7. PS, Ponceau red.

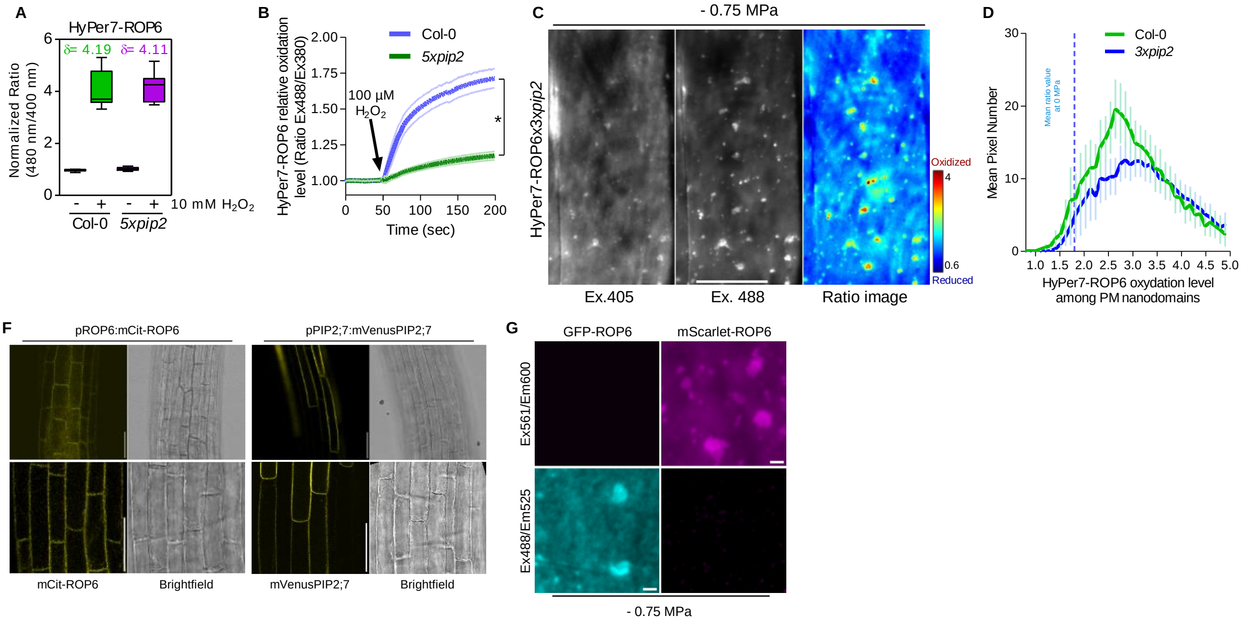


**Figure S4: Characterization of $H_2O_2$ diffusion, osmotically-induced ROS nano-environments.**

A, Relative oxidation dynamics of HyPer7-ROP6 in Col-0 and *5xpip2* backgrounds in response to application of 10 mM exogenous $H_2O_2$. B, Time course ratiometric imaging of 5 days old plantlets expressing HyPer7-ROP6 in Col-0 and *5xpip2* background perfused with 100 µM of exogenous $H_2O_2$. C, Intensity-based (Ex405 and Ex488) and ratio-based (Ratio) TIRF images of HyPer7 expressing root epidermal cells in *3xpip2* after osmotic stimulation. D, Distribution of the pixel counts within HyPer7-ROP6 nanodomains as a function of their ratio values in Col-0 or *3xpip2* after osmotic stimulation. E, Comparison of HyPer7-ROP6 ratio values in Col-0, *3xpip2* or *5xpip2* after osmotic stimulation (-0.75 MPa). F, pROP6:mCit-ROP6 or pPIP2;7:mVenus-PIP2;7 confocal images in root epidermal cells. G, TIRF images root cell expressing either p35S:GFP-ROP6 or pPIN2:mScarlet-ROP6. Images were taken with similar settings.

*Mean with error bars correspond to the 95% confident interval. For B, *is t-test, p-value$_{t=200\ sec}$ =0,0002, n > 16 seedlings from 2 independent replicates. For E, letters show significant differences according to a one-way ANOVA and Tukey's multiple comparisons test at p<0.05. For E , n cells > 20 from n > 15 seedlings from 3 independent replicates. For C, scale bar is 10 µm. For F, scale bar is 50 µm. For G, scale bar is 1 µm*
